# Improved detection and consistency of RNA-interacting proteomes using DIA SILAC

**DOI:** 10.1101/2023.05.18.541276

**Authors:** Thomas Tan, Christos Spanos, David Tollervey

## Abstract

The RNA-interacting proteome is commonly characterized by UV-crosslinking followed by RNA purification, with protein recovery quantified using SILAC labeling followed by data-dependent acquisition (DDA) of proteomic data. However, the low efficiency of UV-crosslinking, combined with limited sensitivity of the DDA approach often restricts detection to relatively abundant proteins, necessitating multiple mass spec injections of fractionated peptides for each biological sample. Here we report an application of data-independent acquisition (DIA) with SILAC in a total RNA-associated protein purification (TRAPP) UV-crosslinking experiment. This gave 15% greater protein detection and lower inter-replicate variation relative to the same biological materials analyzed using DDA, while allowing single-shot analysis of the sample. As proof of concept, we determined the effects of arsenite treatment on the RNA-bound proteome of HEK293T cells. The DIA dataset yielded similar GO term enrichment for RNA-binding proteins involved in cellular stress responses to the DDA dataset while detecting extra proteins unseen by DDA. Overall, the DIA SILAC approach improved detection of proteins over conventional DDA SILAC for generating RNA-interactome datasets, at a lower cost due to reduced machine time.

## Introduction

Isolation and quantitation of the RNA-bound proteome (RBPome) in cells can reveal major regulatory events in RNA function and metabolism that cannot be inferred from total proteomic data. For example, specific ribosomal proteins and translation initiation factors were found to dissociate from mRNA following after glucose withdrawal in yeast, while the cellular abundance of these proteins remained constant (1). UV irradiation has long been used to covalently crosslink RNA with associated proteins. RNA can then be purified, either by affinity (e.g. poly(A) selection) or by enrichment of the total RNA population, and bound proteins characterized. A limitation with all such techniques is the relatively low efficiency of crosslinking. With unmodified RNA, crosslinking efficiency is typically less than 5% (often much less). Incorporating photoactivated nucleotides (e.g. 4-thiouracil) increases the efficiency at each site, but only a fraction of nucleotides are substituted so overall levels are similar. In total RNA-associated proteome purification (TRAPP) a silica column is used to immobilize all RNAs from a denatured cell lysate, allowing RNA-associated proteins to be recovered and identified by mass spectrometry (MS) (1,2).

Most proteomic datasets are currently generated using tandem MS to identify and quantify proteins based on the mass/charge ratio of peptides (MS1) and their fragments (MS2). MS data are typically acquired using either data dependent acquisition (DDA), the traditional acquisition method, or data independent acquisition (DIA) (3). In DDA, individual peptide ions from the MS1 cycle are selected for subsequent MS2 analysis, based on pre-set criteria. The technique is so named because data from MS1 is used to determine what data will be acquired in MS2. Generally, the most intense ions are selected, but this is usually not under control of the operator. This approach has several limitations, including the preferential detection of high-abundance over low-abundance ions, which leads to bias and a lack of sensitivity. In addition, the stochastic element in selecting MS1 features for MS2 analysis results in inconsistent identification of proteins across multiple samples.

In contrast, in DIA all peptides within a defined m/z range are fragmented. This results in a more comprehensive and consistent analysis of the sample, including low-abundance ions that may be missed by DDA. However, the highly complex fragment mass spectra generated from simultaneous fragmentation of multiple precursor ions are challenging to analyze. Therefore, to facilitate the matching of observed and theoretical mass spectra, a spectral library is created that serves as a reference of the observable features in a sample.

Stable isotope labelling by amino acids in cell culture (SILAC) enables the relative quantification of proteins in two or more samples (4). SILAC involves growing cells in media containing different stable-isotope labelled amino acids, such as heavy (^13^C_6_ and/or ^15^N_2_) lysine or arginine, which are incorporated into newly synthesized proteins. The labelled and unlabeled cells are then mixed, lysed, and subjected to MS analysis. The relative abundances of proteins in each sample are determined based on recovery of heavy and light peptides, which are chemically similar but separate in MS. Currently, SILAC data are predominately acquired with DDA; peptide precursors are identified in MS2 whereas quantitation and FDR control are at the MS1 level, which is not always accurate at the resolution of the individual precursor. With recent developments in both mass spectrometers and analysis software, it has become feasible to process SILAC samples by DIA (5–7). A previous study (5) analyzed pulse-SILAC datasets, using a spectral library constructed using narrow-window gas-phase fractionation of the light peptidome. This was followed by *in silico* generation of the heavy spectrum, based on the light spectrum, and single-injection acquisition for experimental samples. Analysis at the proteome scale was not possible with the reported workflow (5). However, proteomic scale SILAC spectral library construction can now be achieved following advances in analysis software, like DIA-NN (8). The library can also be generated entirely *in silico*, using a reference set of FASTA protein sequences. In these approaches the quantitation and quality control of peptide precursors are based on MS2 intensities, and should therefore be accurate for each individual precursor.

In TRAPP, and many other datasets generated following affinity purification, there are no reliable internal reference proteins that allow absolute quantitation of proteins. Previous analyses therefore relied on relative quantitation of proteins using DDA-SILAC. To enhance sensitivity, peptides were reverse phase fractionated using an acetonitrile gradient prior to MS injection, necessitating multiple MS injections for each sample.

Here we report the application of a DIA-SILAC workflow following a modified mammalian TRAPP procedure (9). Data for the effects of arsenite treatment on human RBPome were previously reported (10), so we adopted this as a test system. TRAPP was applied after acute (10 min) and prolonged (1 h) arsenite treatment in human embryonic kidney HEK293T cells with three biological replicates. We aimed to improve sensitivity and accuracy of detection, while eliminating the requirement for pre-fractionation of peptides, thereby enhancing dataset qualities while improving workload and machine time economy.

## Materials and Methods

### Cells

HEK293T cells were grown in DMEM with GlutaMax (Gibco; cat. 10566016) supplemented with 10% fetal calf serum (Sigma; cat. F2442). All cell cultures were maintained at 37°C with 5% CO_2_. Cells were seeded at a confluency of 10% and grown to 70 - 90% before being split or used for experiments. To split cultures, cells were washed once with sterile PBS, then incubated with 0.25% Trypsin-EDTA (Gibco; cat. 25200056; 0.2 times the volume of culture media removed) until detached. Trypsin was inactivated with 5 volumes of media, and cells pelleted at 300 rcf for 5 minutes before resuspension in culture media.

For SILAC experiments, cells were grown for 10 divisions in SILAC DMEM (Thermo Fischer; cat. A33822) supplemented with 10% dialyzed FBS (Sigma-Aldrich; cat. F0392) and 50 µg/L each of lysine and arginine. For “light” cultures, these amino acids were obtained from Sigma. For “heavy” cultures, ^13^C_6_-lysine and ^13^C_6_-arginine were obtained from Cambridge Isotope Laboratories (cat. CLM-226 and CLM-2247).

Sodium arsenite (Merck; cat. 1.06277) was added to the light culture to a final concentration of 1mM and incubated for 10 min or 1 h. An equal volume of water was added to the heavy culture and incubated for the same duration. At the end of the incubation cells were washed with PBS and snap-frozen on the dish in a bath of dry ice/methanol. Cells were stored at - 80°C until crosslinking (still frozen) with 400 mJ/cm^2^ UVC using a Stratalinker 2400 (Stratagene).

### Isolation of the RNA interactome using TRAPP

Matching SILAC pairs were resuspended and homogenized in 3.6 mL of a 1:1 mix between phenol pH 8.0 (Sigma-Aldrich; cat. P4557) and GTC lysis buffer (4 M guanidine thiocyanate, 50 mM Tris-HCl pH 8.0, 10 mM EDTA, and 1% b-mercaptoethanol), and combined in equal proportion (244 µg RNA total in 4 mL phenol-GTC). 0.1 x volume of 3M sodium acetate pH 4.0 was added to the mixed extract and incubated at 80°C for 5 minutes, with 1000 rpm agitation. The mixture was spun at 13,000 rcf for 10 min at RT and supernatant was transferred to a new tube, to which 0.4286 x volume of 100% ethanol was slowly added and mixed by vortexing. The extract was loaded onto three EconoSpin columns for RNA (Epoch Life Science; cat. 1940-250) and each column washed once with 600 µL WB1A (3 M guanidine thiocynate, 1 M sodium acetate pH 4.0, 30% ethanol). Collection tubes were replaced, and the columns were washed once with 600 µL WB1B (2 M guanidine hydrochloride, 20 mM Tris-HCl pH 6.4, 50% acetone, 20% ethanol), twice with 600 µL WB2B (20 mM Tris-HCl pH 6.8, 0.5 mM EDTA, 80% ethanol) and then once with 500 µL WB3B-80 (80% acetonitrile, 50 mM Tris-HCl pH 7.8, 2 mM CaCl_2_). After the last wash columns were spun for 2 min at 16,000 rcf. Collection tubes were replaced with 1.5 mL low protein binding tubes (Thermo Scientific; cat. 90410). To each column was added 0.25 µg Trypsin+Lys-C (Promega; cat. V5073) in 75 µL WB3B-60 (60% acetonitrile, 50 mM Tris-HCl pH 7.8, 2 mM CaCl_2_) with incubation for 16 h at 37°C. The three columns were eluted by spinning at 16,000 rcf for 30 sec, and each column was washed with 75 µL WB3B-60. Eluates from the three columns were pooled and spun at 16,000 rcf for 5 min. Supernatant was transferred to a new tube. 1M DTT (Sigma-Aldrich; cat. 43816) was added to the eluate to a final concentration of 5mM and the mixture was incubated at 37°C for 20 min. Peptide alkylation was done with 10 mM final Iodoacetamide (Merck; cat. A3221) at RT in the dark for 20 min, followed by inactivation with 10 mM final DTT, incubated at RT for 5 min. 0.15 µg Trypsin+Lys-C in 7.5 µL WB3B-60 was added and incubated at 37°C for 1 h for complete digestion of the peptides. The volume of the peptides was evaporated to ∼10 µL and resuspended in 200 µL Buffer A (0.1% NH_4_OH). The material was split at this stage; half was high pH reverse phase fractionated using an SDB-XC stage tip with 10%, 20% and 80% acetonitrile (flow-through was acidified, cleaned with a C18 stage tip and pooled with the 80% fraction) and used for DDA; another half was acidified with 10% TFA to pH ∼1.0 and cleaned with a C18 stage tip and used for DIA without fractionation.

### Mass spectrometry

For library generation, 30 µg of total protein from HEK293T cells grown in Light medium were digested using the Filter Aided Sample Preparation (FASP) protocol (11) with minor modifications. In brief, protein samples were added on top of 30 kDa MWCO filter units (Vivacon, UK), along with 150 µl of denaturation buffer (8M Urea in 50mM ammonium bicarbonate (ABC) (Sigma Aldrich)) and spun at 14,000 x g for 20 min. An additional wash with 200 µl of denaturation buffer was performed under the same conditions. The protein sample was then reduced with 100 µl of 10 mM dithiothreitol (Sigma Aldrich, UK) in denaturation buffer for 30 min at ambient temperature, and alkylated by adding 100 µl of 55 mM iodoacetamide (Sigma Aldrich, UK) in denaturation buffer and incubated for 20 min at ambient temperature in the dark. Two washes with 100 µl of denaturation buffer and two with digestion buffer (50mM ABC) were performed under the conditions described above, before the addition of trypsin/LysC mix (Pierce, UK). The protease:protein weight ratio was 1:50 and proteins were digested overnight at 37°C. Following digestion, the sample was spun at 14,000 x g for 20 min and the flow-through containing digested peptides was collected. Filters were rinsed with 100 µl of digestion buffer and the flow-through collected. Eluates from the filter units were acidified using 20 µl of 10% Trifluoroacetic Acid (TFA) (Sigma Aldrich), and spun onto C_18_ StageTips as described (12). Peptides were eluted in 40 μL of 80% acetonitrile in 0.1% TFA and concentrated down to 1 μL by vacuum centrifugation (Concentrator 5301, Eppendorf, UK). Peptide samples were then prepared for LC-MS/MS analysis by dilution to 5 μL using 0.1% TFA.

LC-MS analyses were performed on an Orbitrap Exploris™ 480 Mass Spectrometer (Thermo Fisher Scientific, UK), coupled on-line to an Ultimate 3000 HPLC (Dionex, Thermo Fisher Scientific, UK). Four 2µg-injections of the HEK cell digest were performed. Peptides were separated on a 50 cm (2 µm particle size) EASY-Spray column (Thermo Scientific, UK), which was assembled on an EASY-Spray source (Thermo Scientific, UK) and operated constantly at 50°C. Mobile phase A consisted of 0.1% formic acid in LC-MS grade water and mobile phase B consisted of 80% acetonitrile and 0.1% formic acid. Peptides were loaded onto the column at a flow rate of 0.3 μL min^-1^ and eluted at a flow rate of 0.25 μL min^-1^ according to the following gradient: 2 to 40% mobile phase B in 120 min and then to 95% in 11 min. Mobile phase B was retained at 95% for 5 min, then returned to 2% in a minute, and stayed at 2% until the end of the run (160 min).

For gas phase fractionation, survey scans were recorded at 120,000 resolution (scan range 350-1650 m/z) with an ion target of 5.0E6, and injection time of 20 ms. MS2 DIA was performed in the orbitrap at 30,000 resolution, maximum injection time of 55 ms and AGC target of 3.0E6 ions. We used HCD fragmentation (13) with stepped collision energy of 25.5, 27 and 30. For the first injection the scan range was between 400 and 600 m/z with isolation windows of 5 m/z. For the other three injections the scan range was 600-800, 800-1000 and 1000-1400, with 5 m/z isolation windows and default charge state of 3.

For SILAC TRAPP samples acquired with DDA, the LC-MS preparation and the injection conditions were as described above. MS1 was performed at 120,000 resolution with scan range 350-1500 m/z, AGC target of 3.0E6 ions and injection time of 50 ms. MS2 scans were performed at resolution of 15,000 with isolation window of 1.4 Thompson AGC target of 8.0E5 and injection time of 60 ms.

For SILAC TRAPP samples acquired with DIA, LC-MS conditions were the same as above. Variable isolation windows was used as the table below:

| m/z | z | t start (min) | t stop (min) | Isolation Window (m/z) |
| --- | --- | --- | --- | --- |
| 383.375 | 3 | 0 | 185 | 66.8 |
| 423 | 3 | 0 | 185 | 13.5 |
| 435 | 3 | 0 | 185 | 11.5 |
| 446.5 | 3 | 0 | 185 | 12.5 |
| 458 | 3 | 0 | 185 | 11.5 |
| 469 | 3 | 0 | 185 | 11.5 |
| 480 | 3 | 0 | 185 | 10.5 |
| 490.5 | 3 | 0 | 185 | 10.5 |
| 501 | 3 | 0 | 185 | 11.5 |
| 512 | 3 | 0 | 185 | 11.5 |
| 523 | 3 | 0 | 185 | 11.5 |
| 533.5 | 3 | 0 | 185 | 10.5 |
| 544 | 3 | 0 | 185 | 11.5 |
| 554.5 | 3 | 0 | 185 | 10.5 |
| 565 | 3 | 0 | 185 | 11.5 |
| 575.5 | 3 | 0 | 185 | 10.5 |
| 586 | 3 | 0 | 185 | 11.5 |
| 597.5 | 3 | 0 | 185 | 12.5 |
| 609.5 | 3 | 0 | 185 | 12.5 |
| 621.5 | 3 | 0 | 185 | 12.5 |
| 633 | 3 | 0 | 185 | 12.5 |
| 645 | 3 | 0 | 185 | 13.5 |
| 657.5 | 3 | 0 | 185 | 12.5 |
| 670.5 | 3 | 0 | 185 | 14.5 |
| 684 | 3 | 0 | 185 | 13.5 |
| 697 | 3 | 0 | 185 | 13.5 |
| 710.5 | 3 | 0 | 185 | 14.5 |
| 725.5 | 3 | 0 | 185 | 16.5 |
| 741 | 3 | 0 | 185 | 15.5 |
| 756.5 | 3 | 0 | 185 | 16.5 |
| 773.5 | 3 | 0 | 185 | 18.5 |
| 791 | 3 | 0 | 185 | 17.5 |
| 808.5 | 3 | 0 | 185 | 18.5 |
| 827 | 3 | 0 | 185 | 19.5 |
| 846.5 | 3 | 0 | 185 | 20.5 |
| 866.5 | 3 | 0 | 185 | 20.5 |
| 887.5 | 3 | 0 | 185 | 22.5 |
| 910.5 | 3 | 0 | 185 | 24.5 |
| 935.5 | 3 | 0 | 185 | 26.5 |
| 962.5 | 3 | 0 | 185 | 28.5 |
| 992 | 3 | 0 | 185 | 31.5 |
| 1025 | 3 | 0 | 185 | 35.5 |
| 1063 | 3 | 0 | 185 | 41.5 |
| 1108.5 | 3 | 0 | 185 | 50.5 |
| 1391.62 | 3 | 0 | 185 | 516.8 |

For DDA-SILAC samples, MaxQuant software platform (14) version 1.6.1.0 was used to process the raw files and the search was conducted against the *Homo sapiens* protein database (released 14/05/19), using the Andromeda search engine (15). For the first search, peptide tolerance was set to 20 ppm while for the main search peptide tolerance was set to 4.5 ppm. Isotope mass tolerance was 2 ppm and maximum charge set to 5. Digestion mode was set to specific with trypsin and LysC allowing maximum of two missed cleavages. Carbamidomethylation of cysteine was set as fixed modification. Oxidation of methionine was set as variable modification. Multiplicity was set to 2, specifying Arg6 and Lys6 for heavy labels. Peptide and protein identifications were filtered to 1% FDR.

To process the DIA raw files the DIA-NN software platform (8) version 1.8.1 was used, searching the Uniprot one protein sequence per gene database (Proteome ID UP000005640, released in September, 2022). The *in silico* spectral library was constructed with the predicted light library, using parameters and modifications as previously described (6). The project-specific library was built using gas phase fractionated data, described above. Precursor ion generation was based on the chosen protein database with deep-learning based spectra, retention time and IMs prediction. Regarding general parameters, digestion mode was set to specific with trypsin, allowing maximum of one missed cleavage. Carbamidomethylation of cysteine was set as fixed modification. Oxidation of methionine, and acetylation of the N-terminus were set as variable modifications. The parameters for peptide length range, precursor charge range, precursor m/z range and fragment ion m/z range as well as other software parameters were used with their default values. The precursor FDR was set to 1%, and then further filtered using 1% Translated Q value.

### DDA data processing

For non-crosslinked vs crosslinked comparisons, raw intensities of light and heavy samples in each SILAC pair were obtained from the proteinGroups.txt file generated by MaxQuant. Intensities were normalized by mean using NormalyzerDE version 1.18.0 in R version 4.3.0. Statistical tests were performed using 2-sided T Tests with Benjamini Hochberg multiple test correction.

For arsenite-treated vs mock comparisons with MaxQuant normalized datasets, normalized Heavy/Light ratios for each protein were obtained from the MaxQuant proteinGroups.txt file. Heavy/Light ratios were inverted to Light/Heavy. For Vsn and Cyclic Loess normalized datasets, Heavy and Light intensities for each peptide were obtained from the peptides.txt file; only peptides labelled as unique were used. Peptide intensities were normalized using NormalyzerDE, with 10 min light as Group1, 10 min heavy as Group2, 1 h light as Group3 and 1 h heavy as Group4. Light/Heavy ratio for each peptide in the normalized datasets were calculated; missing values were replaced with “NA”. For every gene in each sample, the number of non-NA values was calculated; if the number was 1, the non-NA value was replaced by “NA”; otherwise the value was retained. Median values for all peptides assigned to each gene were calculated and used to represent the level of the protein in the sample.

Proteins annotated as contaminants, reverse, or keratins were removed. Only proteins that have non-NA SILAC ratios in 2 out of 3 replicates at any time point were further analyzed. Proteins analyzed were restricted to those represented by 2 or more peptides and detected in at least 2 out of 3 replicates, at both time points in non-treated controls (SILAC heavy). At this point the datasets were named the “processed datasets”.

Differential expression analysis was performed on proteins that have non-NA Light/Heavy ratios in 2 out of 3 replicates at both time points, using 2-sided T Test with Benjamini Hochberg multiple test correction. Significantly changed proteins were defined as those with T Test P-value<0.05 and FC>90^th^ percentile and <10^th^ percentile. Proteins deemed significant after BH correction (FDR<0.05) were separately marked. GO term enrichment analysis was done on WebGestalt (16) using Molecular Function noRedundant database, with a list of all detected proteins in the DDA dataset as reference.

### DIA data processing

Intensities for each peptide were obtained from the “Precursor.Translated” column of the Report.tsv generated by DIA-NN. Precursors with Precursor.Id containing (SILAC-K-H) and/or (SILAC-R-H) were labelled as heavy and those containing (SILAC-K-L) and/or (SILAC-R-L) were labelled as light. Samples to which each peptide belonged were determined by the “Run” column of Report.tsv and concatenated with either H or L depending on whether the peptide is heavy or light. “Protein.Ids”, “Genes”, and “Precursor.Id” with (SILAC-K-L/H) and (SILAC-R-L/H) removed were concatenated to form the identity of each peptide. Together a long-formatted file containing 3 columns: Sample, Precursor, Intensity was formed. A matrix file was generated using Pivot Table in Excel with Sample as columns, Peptide as rows, and Intensity as values. The concatenated Precursor column was separated again. This matrix file was used as the input file for NormalyzerDE with 10 min light as Group1, 10 min heavy as Group2, 1 h light as Group3 and 1 h heavy as Group4. Datasets normalized by Median, Vsn, and Cyclic Loess were analyzed.

Light/Heavy ratios for each peptide in each sample were calculated, with missing values replaced with “NA”. For every gene in each sample, the number of non-NAs was calculated; if the number was 1, the non-NA value was replaced by “NA”; otherwise the value was retained. Median ratios of all peptides belonging to a gene was calculated and used to represent the ratio of the protein in the sample. Only proteins that have non-NA SILAC ratios in 2 out of 3 replicates at any time point were further analyzed. Proteins for analysis were restricted to those represented by equal or more than 2 peptides and detected in at least 2 out of 3 replicates in both time points in non-treated controls (SILAC heavy). At this point the datasets were named the “processed datasets”.

Differential expression analysis was performed on proteins that have non-NA Light/Heavy ratios in 2 out of 3 replicates at both time points, using 2-sided T Tests with Benjamini Hochberg multiple test correction. Significantly changed proteins were defined as those with T Test P-value<0.05 and FC>90^th^ percentile and <10^th^ percentile. Proteins deemed significant after BH correction (FDR<0.05) were separately marked. GO term enrichment analysis was done on WebGestalt using Molecular Function noRedundant database and a list of all detected proteins in the DIA dataset as reference.

### Data imputation

The effect of imputation by Random Forest (using R 4.2.2 with missForest library version 1.5) and MIN*0.5 (replacement of missing values by 0.5 x the minimum value of the sample) on the normalized datasets (MaxQuant normalized for DDA and Cyclic Loess normalized for DIA) was analyzed for all proteins detected in non-treatment controls, and proteins enriched by UV-crosslinking and significantly changed (according to DIA) between 10 min and 1 h of treatment.

## Results

DIA detects more RNA-binding proteins (RBPs) with lower inter-replicate variance. For mammalian cell SILAC-TRAPP cells were pre-grown for 10 divisions, in medium containing either standard “light” lysine and arginine or “heavy” ^13^C_6_-lysine and ^13^C_6_-arginine. At T0, light cultures were treated with arsentite (1mM final), while heavy cultures were mock treated. Cells were snap-frozen after 10 min or 1 h, then UV-irradiated at -80°C and kept frozen until used. Since some proteins, such as glycoproteins, bind to silica column with high affinity irrespectively of their binding to RNA, four non-crosslinked non-treated samples (two light and two heavy) at T0 were compared with four crosslinked non-treated samples at the same time point using DDA. Proteins were regarded as truly RNA-binding if detected only in crosslinked samples, or significantly enriched (Two tailed T test with Benjamini Hochberg correction, FDR<0.05, FC>1.5, using mean-normalized protein intensities) (Supplementary data 1).

For SILAC-TRAPP analysis, frozen cells were directly resuspended in cell lysis buffer. Total cell lysate containing 122 µg of RNA from each light sample was mixed an equal amount of lysate from the heavy SILAC pair. RNA complexes were isolated on silica columns and the co-purified proteins were recovered from the column by Trypsin plus Lys-C digestion. Released peptides were separated for either DDA or DIA acquisitions on an Orbitrap Exploris™ 480 Mass Spectrometer (procedures detailed in Material and Methods). To achieve good detection, DDA MS required injection of three fractions from an acetonitrile gradient per biological sample. In contrast, DIA samples were injected as single-shot. For project-specific spectral library generation, four injections were used (Fig. 1A), but this can be omitted as described below.

**Fig. 1.**
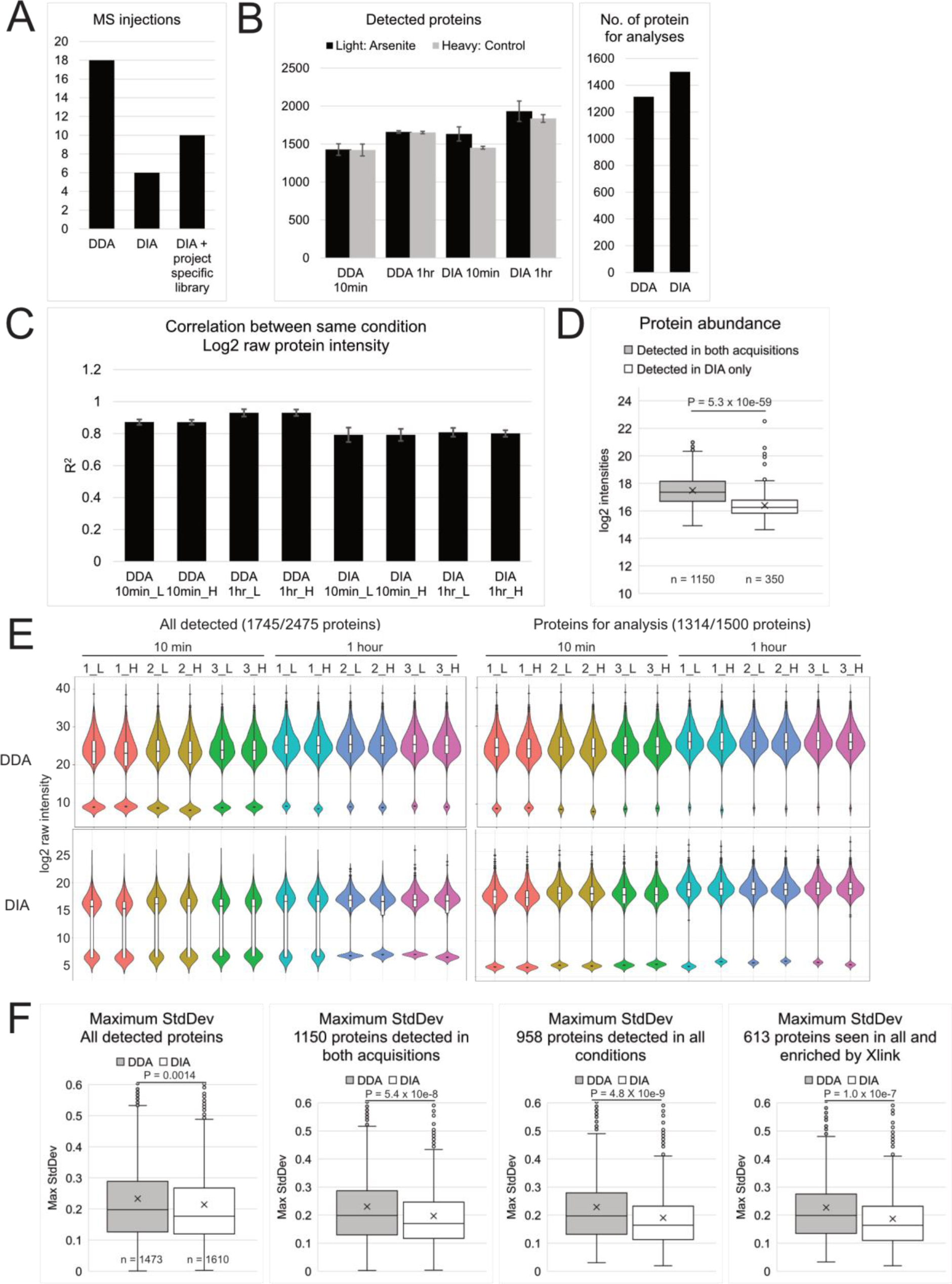
DIA detected more proteins with lower inter-replicate variability compared to DDA. (A) Left: Number of mass-spec injections required by the two acquisitions. (B) Left: means (bar) and standard deviations (error bar) of number of detected proteins in 3 technical replicates were plotted for all 4 conditions; Right: number of proteins for analysis, defined as those represented by more than 2 peptides, and detected in at least 2 out of 3 control samples (heavy) in both time points. (C) Pearson correlation coefficient (R^2^) between log2 raw protein intensities in 3 replicates of the same conditions. Bar, mean; error bar, standard deviation. (D) Box plot showing average log2 intensities of proteins detected in both acquisitions, and proteins detected only in the DIA dataset. (E) Violin and box plot showing log2 raw intensity of all detected proteins or proteins for analysis. Colors denote the SILAC pairs. Missing values were imputed with half of the minimal log2 value of each sample, reflected by the second population below the main ones in each violin plot. (F) Box plot showing maximum standard deviation (higher of the two time points) of light/heavy ratios of the 3 replicates for all detected proteins, proteins detected in both acquisitions, proteins detected in all conditions, and proteins detected in all conditions and enriched in UVC-crosslinked over non-crosslinked samples. Center line, median; cross, mean; box limits, upper and lower quartiles; whiskers, 1.5x interquartile range; points, outliers. P values were determined by 2-sided T Tests, with n numbers indicated in each graph.

DDA datasets were processed with MaxQuant version 1.6.1.0 (14). DIA datasets were processed with DIA-NN version 1.8.1 (8) with *in silico* spectral library constructed from the reference human, one sequence per gene, proteome (Uniprot ID UP000005640) (17). A protein was considered detected if represented by 2 or more quantitated peptides, otherwise the protein quantity was replaced with NA for that sample. For DDA datasets, peptide quantity normalization and protein quantity calculation were performed using MaxQuant. For DIA datasets, peptide quantities were normalized with NormalyzerDE (18) using Cyclic Loess. Protein quantity was calculated as the median abundance of all represented peptides, using absolute quantities in Figure 1 and SILAC ratios thereafter.

To assess the relative sensitivity of DDA and DIA acquisition, we first determined total numbers of detected proteins, initially defined as all proteins with one or more intensity in any condition (Fig. 1B left). The DIA datasets had more detected proteins (mean = 1,542 in 10 min samples; 1,885 in 1 h samples, decimal point rounded up) than corresponding DDA datasets (1,425 at 10 min; 1,656 proteins at 1 h). Variations between replicate samples were low, indicating consistent sensitivity. In DIA datasets, more proteins were detected in the arsenite-treated samples compared to control at both 10 min (1,633 vs 1,451) and 1 h (1,932 vs 1,837), while numbers of detected proteins were similar in DDA datasets (1,428 vs 1,421 at 10 min, 1,660 vs 1,652 at 1 h). Since non-treated controls at 10 min and 1 h should be biologically identical, discrepancies in numbers of detected proteins likely reflect variations in sample preparation. To account for this variability, downstream analyses were restricted to proteins detected in at least 2 out of 3 replicates in the untreated control at any time point (Fig. 1B right, 1,314 proteins for DDA and 1,500 proteins for DIA). Since the downstream analysis is restricted to proteins consistently detected in the untreated control, any proteins with missing light:heavy ratios will reflect the effects of arsenite on the treated samples.

Variations in quantitation between replicates in the same conditions were quantified by the correlation coefficient (R^2^) of log2 raw (not normalized) intensities of proteins detected in any two of the three heavy replicates (Fig. 1C). Raw DDA dataset had higher correlation coefficients (mean R^2^=0.901) and consistency was good between correlations (mean SD=0.0189) compared to raw DIA (mean R^2^=0.798, SD=0.0327), reflecting a higher inter-replicate variance in the DIA dataset, possibly due to lower quantitation precisions for the extra proteins detected.

Proteins detected exclusively in the DIA dataset had lower abundances compared to those detected in both acquisitions (Fig. 1D), with a log2 intensity median of 16.25 compared to 17.36, 2-tailed T Test P-value 5.34 x 10^-59^. Distribution of raw protein intensities were displayed as violin and box plots (Fig. 1E), with missing values imputed with 0.5 x minimum value of the sample. 10 min samples in both acquisitions had more missing values for all detected proteins (mean of 18.4% in DDA and 37.7% in DIA) than 1 h samples (5.1% in DDA and 23.9% in DIA). This was reduced for proteins passing the threshold for analysis (detected in at least 2 out of 3 replicates in the heavy samples), with 8.16% missing values in DDA and 23.1% in DIA at 10 min, and 1.35% in DDA and 12.3% in DIA at 1 h. DDA data were also more widely distributed, with an average log2 interquartile range of 4.33 compared to 1.88 in DIA. Maximum inter-replicate standard deviations of Light/Heavy ratios for individual proteins in DDA and DIA were also compared (Fig. 1F). The median and mean maxima SD in DIA were consistently lower than those in DDA, with increasing divergence when restricted to proteins detected in both acquisitions (DDA and DIA), in all conditions (10 min and 1 h), and in all conditions with enrichment following UV-crosslinking (protein list presented in Supplementary data 1). In the most restricted protein list, DDA dataset had a median/mean maximum SD of 0.20/0.23 while DIA had MaxSDs of 0.16/0.19, indicating a better inter-replicate consistency in DIA compared to DDA for these proteins.

### Comparing quantitation by DDA and DIA for RBP recovery

Several dataset normalization approaches were implemented in MaxQuant and NormalyzerDE; applying MaxQuant, Vsn and Cyclic Loess algorithms for DDA and Median, Vsn and Cyclic Loess for DIA. Table 1A shows correlations between the two datasets using different combinations of normalization methods. MaxQuant normalization was chosen for DDA and Cyclic Loess for DIA. A more detailed comparison between the algorithms is described below.

**Table 1.** Correlation between, and level of inter-replicate variation in DDA and DIA datasets normalized with MaxQuant/Median, Vsn, or Cyclic Loess. (A) Pearson correlation coefficient (R^2^) values for log2FC of individual proteins between datasets normalized by different methods, calculated using proteins detected in all conditions. (B) Median and mean of maximum standard deviation between replicates for proteins detected in all conditions and enriched by crosslink, in datasets normalized with different algorithms.

| DDA \ DIA | Median | Vsn | CycLoess |
| --- | --- | --- | --- |
| MaxQuant | 0.514 | 0.515 | 0.512 |
| Vsn | 0.480 | 0.482 | 0.476 |
| CycLoess | 0.468 | 0.470 | 0.470 |

**Table 1. Correlation between, and level of inter-replicate variation in DDA and DIA datasets normalized with MaxQuant/Median, Vsn, or Cyclic Loess.**
| Max SD | Median | Mean |
| --- | --- | --- |
| DDA-MaxQuant | 0.199 | 0.227 |
| DDA-Vsn | 0.205 | 0.249 |
| DDA-CycLoess | 0.204 | 0.247 |
| DIA-Median | 0.201 | 0.220 |
| DIA-Vsn | 0.163 | 0.189 |
| DIA-CycLoess | 0.164 | 0.187 |

Distributions of Light/Heavy ratios across datasets were plotted in Fig. S1. Apart from proteins having missing values at one of the time points (NAs were plotted as 0), the ratios distributed normally in both acquisitions, allowing 2-sided T Tests to be applied for the differential RNA-association analysis between 10 min and 1 h time points. The statistical test was also complemented with Benjamini Hochberg (BH) multiple test correction to minimize false discovery, however as the numbers of significant proteins were low after correction (6 in DDA and 9 in DIA), P-values from the T Test were used as the main selection criterion for proteins with significantly altered RNA association. 10^th^ and 90^th^ percentile of log2 fold-change in average ratio from 10 min to 1 h (log2FC) were used as additional selection criteria. Log2 fold changes of individual proteins in each dataset were also used to assess the similarity between DDA and DIA datasets. Proteins with a P-value <0.05 and a fold-change >90^th^ and <10^th^ percentile of the dataset, were regarded as significantly changed in RNA interaction between the 10 min and 1 h time points. The 10^th^/90^th^ percentile for log2FC was -0.4883/0.2539 in DDA and -0.3913/0.3196 in DIA, these values were used as the significance threshold instead of an arbitrary value (commonly 0.5850 for 1.5-fold or 1 for 2-fold) to maintain a consistent stringency between the datasets.

Both datasets contain proteins which were either completely missing at one time point, or have only one quantitation at any time point (245 in DDA and 271 in DIA, Fig. 2A). Since DDA-DIA correlation for these proteins were very poor (assessed in Fig. S2), they were excluded from the differential RNA association analysis.

**Fig. 2.**
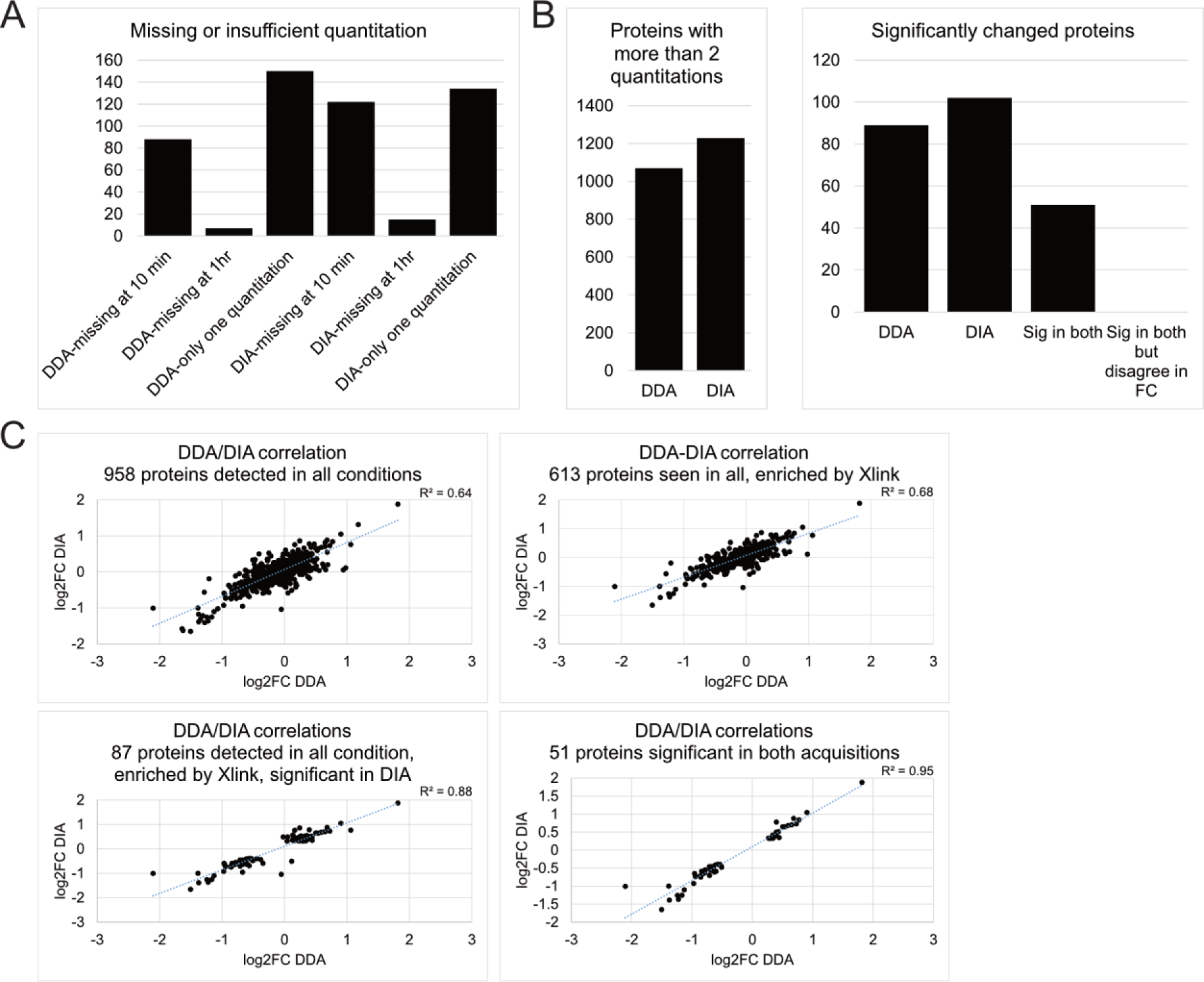
Positive correlation between protein quantitation and fold-change in SILAC light/heavy ratio between time points in the DDA and DIA datasets. (A) Number of proteins excluded from DE analysis, including those missing at 10 min or 1 h, or having only one quantitation at any time points. (B) Left: Number of proteins with at least 2 quantitations at any time point in DDA and DIA datasets. Right: Number of significantly changed proteins (2-sided T Test P-value<0.05, FC>90^th^ and <10^th^ percentile) in each acquisition, common significant proteins in both datasets, proteins significant in both datasets but have opposite fold-changes. (C) Dot plot of log2 fold change in light/heavy ratio from 10 min to 1 h time points in DDA and DIA datasets for proteins detected in all conditions; proteins detected in all conditions and enriched by crosslinking; proteins detected in all conditions, enriched by crosslinking, and significantly changed according to the DIA dataset; and proteins commonly significant in both datasets.

From the remaining proteins with sufficient quantitation (1,069 in DDA and 1,229 in DIA, Fig. 2B left), we identified 89 proteins in DDA and 102 proteins in DIA showing significant changes in RNA association between the 10 min and 1 h time points of arsenite treatment, that were also detected at both time points in the non-treated controls and enriched in UV-crosslinked samples. Of them, 51 proteins were consistently significant in both datasets (Fig. 2B right).

DDA/DIA correlation also improved progressively when proteins for analysis became more restricted, with R^2^ value of 0.64 for proteins detected in all conditions; 0.68 for proteins detected in all conditions while enriched by crosslinking; 0.88 for proteins detected in all conditions, enriched by crosslinking, and significant according to DIA; and 0.95 for crosslinking-enriched proteins significant in both acquisitions (Fig. 2C). Together these results indicate that quantitation in the DIA dataset was similar to the DDA dataset, when potential sources of noises were eliminated. Also, for proteins missing or with insufficient quantitation at one time point, their dynamics are not reliably measured in either acquisition.

### Enrichment for cellular stress response factors in significantly altered RBPs

A well characterized cellular mechanism observed following arsenite exposure is the integrated stress response. Translation of mRNAs into proteins is halted and stress granules are formed to sequester mRNAs for which translation is inhibited. This occurs between 30 min and 1 h of treatment with high arsentite dosages (0.5-2.5 mM) (19,20). Comparing the RNA-interactomes at 10 min and 1 h of arsenite treatment, should therefore reveal increasing recovery of translation regulators and stress granule-related factors. Since the DDA-DIA correlation is good for proteins detected in all conditions (those shown in Fig. 2B), a GO-term enrichment analysis was performed using these proteins with significantly altered RNA-association in the two datasets (Fig. 3 left). Manually selected proteins possibly linked to stress responses are shown in Fig. 3 right. GO terms in dark blue showed significant enrichment (FDR<0.05); terms in light blue were significant by P-value (<0.01) but not FDR (>0.05). Proteins highlighted in bold brown were deemed significant after BH multiple test correction (FDR<0.05).

**Fig. 3.**
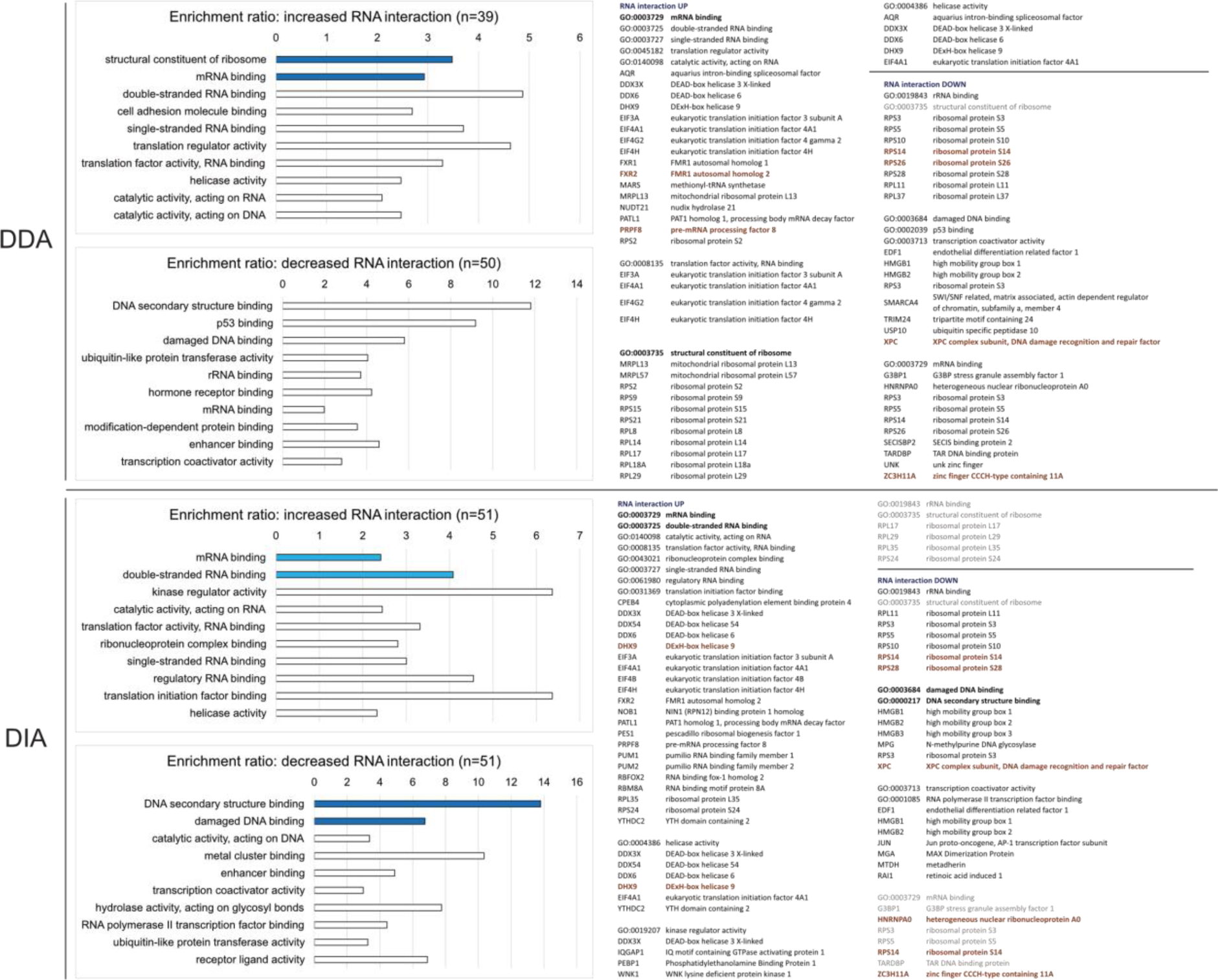
Significantly differentially associated proteins in DDA and DIA datasets have similar GO term enrichment. Left: top-ten enriched GO terms in molecular function for proteins with increased or decreased RNA interaction from 10 min to 1 h and detected at both time points according to the DDA or DIA dataset. Dark blue bar, FDR<0.05; pale blue bar, P-value<0.01 when all FDRs are ≥0.05; white bar, FDR>0.06. GO terms were sorted by FDR in ascending order, or by pre-adjusted P value if all FDRs were 1. Right: selected gene sets possibly related to the stress-induced RNA translation shutdown. Font: bold black, enriched GO term with FDR<0.05 or P-value<0.01 when all FDRs were ≥0.05; black, enriched with FDR>0.05; grey, GO term absent in top-ten enrichment; bold brown, proteins significant according Benjamini-Hochberg multiple test correction, FDR<0.05.

For both DDA and DIA, proteins with significantly increased RNA-interactions from 10 min to 1 h were frequently linked to GO terms related to mRNA binding, single or double-stranded RNA binding and ribonucleoprotein complexes, consistent with the expected regulation in mRNA translation. However, the DIA dataset called one extra DEAD-box helicase DDX54 while the DDA dataset had more ribosomal proteins. Both datasets included translation regulators, including translation initiation factor eIF3A, eIF4A1 and eIF4H, DEAD/DExH-box helicases DDX3X, DDX6 and DHX9, and other stress granule related factors, including G3BP1 and FXR2 (21). For proteins with significantly decreased RNA-interactions, both datasets included specific ribosomal proteins with known extraribosomal functions (22) including RPS3, RPS10, RPS14 and RPS28. These decreases in RNA interactions likely indicate changes in specific mRNA regulation, rather than a global reduction in mRNA translation. Specifically, RPS3 and RPS10 are known to regulate translation of TISU element-containing mRNAs (23), which are enriched in proteins involved in ribosome, translation, and RNA processing (24). RPS28 was shown to regulate expression of specific proteins enriched in the germline or with anti-ageing functions in *Drosophila* (25). Several proteins detected only in DIA were significantly changed in RNA association following prolonged arsenite treatment. These included JUN (down), a crucial component of the AP-1 heterodimer transcription complex (26); Coiled-Coil Domain Containing proteins CCDC9 (up), and CCDC124 (down), which participate in RNA metabolism; cell cycle regulator ATAD5 (down); and putative mRNA translation regulator DRG1 (down). In contrast, all proteins significantly detected in DDA datasets were also detected by DIA. A RAF/MEK complex inhibitor, PEBP1, showed significantly increased RNA association in both DDA and DIA. Inhibition of stress-activated c-Jun N-terminal kinase (JNK) activity by the PEBP1/MEK/JNK/JUN axis potentially mediates the dissociation of JUN from RNA identified above.

Together, proteins significantly changed in the DDA and DIA datasets were similar, with DDA calling more ribosomal proteins and DIA calling more stress-responsive helicases. DIA detected functionally relevant proteins that were missed by DDA, supporting the more comprehensive nature of the DIA dataset.

### Comparing normalization methods for RBP recovery

Previous analyses on DDA/DIA correlation and differential RNA interaction were based on MaxQuant-normalized DDA dataset and Cyclic Loess-normalized DIA dataset. To test whether other methods of normalization enhance DDA/DIA correlations and/or inter-replicate variation, the two datasets were separately normalized by MaxQuant (DDA) and Median (DIA), Vsn, or Cyclic Loess. The Vsn algorithm was chosen due to its reported superior performance in reducing inter-replicate variation in total proteomic datasets (27). Cyclic Loess was originally designed to handle two-color microarrays with different efficiencies of dye incorporation (28), similar to the concept of Light and Heavy incorporation in SILAC experiments. Effects of these algorithms on the distribution of peptide intensities are illustrated as violin and box plots in Fig. S3. Since MaxQuant normalization was performed at the level of Light/Heavy ratio and not on peptide intensities, median-normalized intensities, on which MaxQuant normalization was based (29), was plotted. Correlations between DDA and DIA datasets were measured by the R^2^ value between log2FC of proteins detected in all conditions (Table 1A). Variation between replicates was measured by the maximum standard deviation (greater of the 10 min and 1 h time points) between normalized Light/Heavy ratios (Table 1B). Amongst these methods, the MaxQuant/Vsn pair produced the highest R^2^ value of 0.514, while Median and Cyclic Loess normalization produced similar but lower R^2^ values. Vsn, and similarly Cyclic Loess, reduced the median and mean maximum SD for DIA, but not for the DDA dataset. Together MaxQuant for DDA dataset gave the best DDA-DIA correlation, while Vsn and Cyclic Loess behaved similarly in the DIA dataset.

### Comparing imputation methods for RBP recovery

The differential RNA association analysis above was based on the processed datasets, which included only proteins quantified more than twice at each time point. We also tested whether two imputation methods for handling missing values affect DDA/DIA correlations and significance tests. Based on the assumption that values are missing due to low abundance, we tested the use of 0.5 x the minimum Light/Heavy ratio in the sample (MINx0.5). Alternatively, machine learning can be applied, training the algorithm to predict the missing ones based on the observed values. Random Forest (RF) (30) uses such a strategy and was reported to out-perform other imputation algorithms in LC-MS metabolomic (31) and label-free proteomic data (32) for predicting values missing at random. The two methods were used here to impute both DDA and DIA datasets at the protein level (Light/Heavy ratios of the 3 replicates at 2 time points, normalized using the MaxQuant/Cyclic Loess pair). Proteins quantified at least twice at either time point were included in the analysis. DDA-DIA correlations are shown in Fig. 4A, for proteins detected in both acquisitions, and those significant according to DDA, or DIA. Significant proteins (numbers in Fig. 4B) were selected using the same criteria as previous analyses (2-sided T Test P-value <0.05, FC >90^th^ and <10^th^ percentile) and listed in Table S1 together with their top-ten enriched GO terms in Fig.S4.

**Fig. 4.**
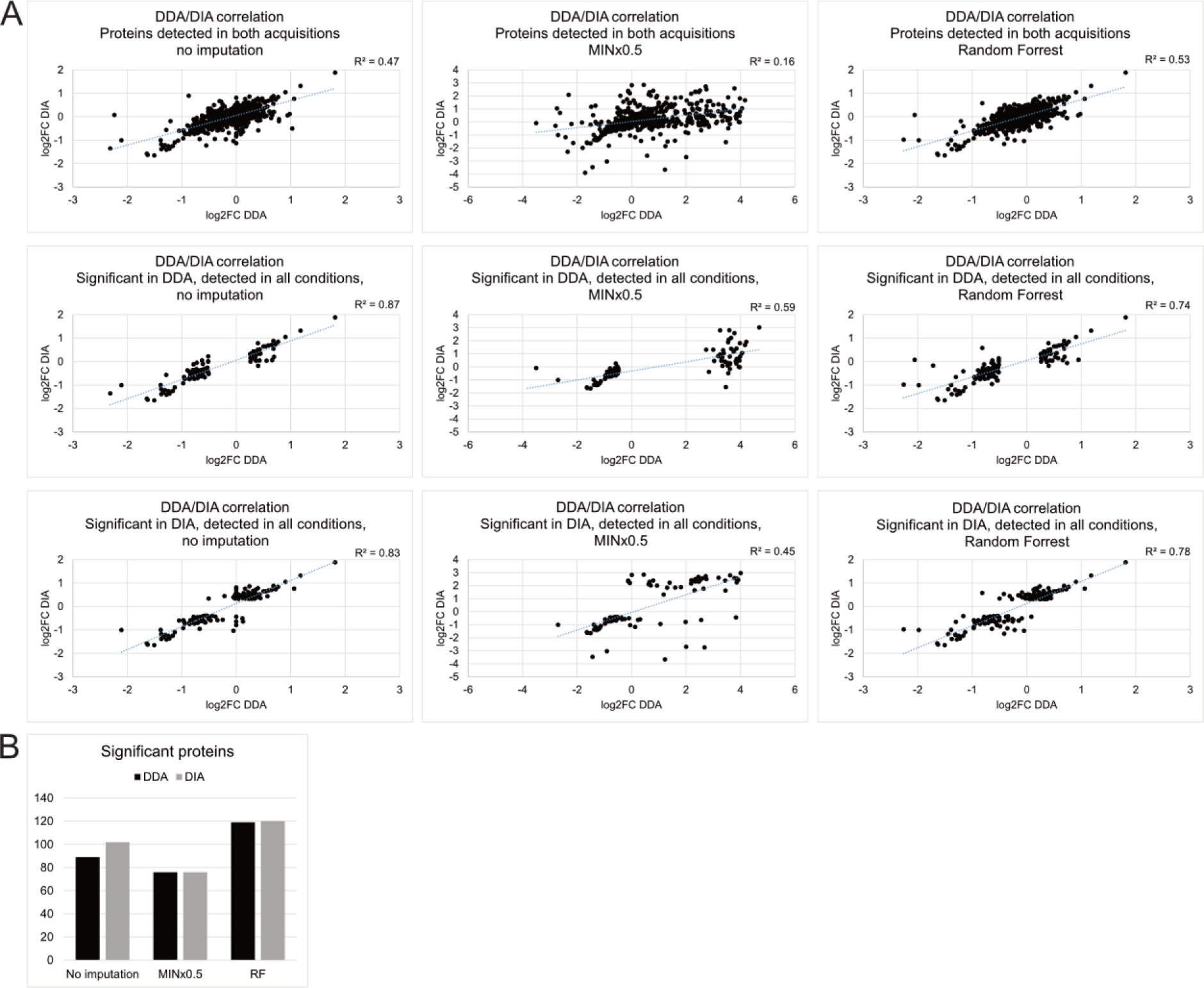
Effects on the datasets by different imputation approaches. Imputation by MINx0.5 replaced significant proteins with those missing at 10 min, while Random Forrest produced similar dataset to no imputation, but called slightly more significant proteins. (A) Like-for-like comparison of DDA/DIA correlation between original datasets with no imputation, with missing values replaced with sample minimum log2 value minus 1 (MINx0.5) or using Random Forrest. Comparison was restricted to proteins detected in both acquisitions and proteins detected in all conditions while significantly changed between 10 min and 1 h of treatment according to DDA or DIA. (B) Number of proteins regarded as significantly changed between 10 min and 1 h of treatment (2-sided T test P-value<0.05, FC >90^th^ and <10^th^ percentile, enriched in UV-crosslinked samples) in datasets with no imputation, imputed by MINx0.5 or Random Forrest.

Since the MINx0.5 approach assumes all missing values were due to low abundance, and most missing values are present at 10 min, the majority of imputed proteins became highly increased in RNA association from 10 min to 1 h. This may be an approach to identify proteins that are associated to RNA only after prolonged arsenite exposure. However this will only be true if the missing values did not occur at random. In the DIA dataset, notable proteins included stress-responsive translation factor EIF2A; RNA quality control factor SUPV3L1; DRG1 stabilizer ZC3H15; splicing regulator MBNL3; 60S ribosomal subunit nuclear export regulator NMD3; ribonuclease P and MRP complex components RRP25L and RRP30; and RAF/MEK complex inhibitor PEBP1, the last one is significant according to all three approaches.

Random Forrest in comparison, retained the overall distribution of the datasets while assigning predicted values to those missing at any time point. Although with only one or no known values at the missing time point to aid prediction, most of these missing proteins did not pass the significance threshold. In the DIA dataset, 5 out of 102 (CDCA2, CNOT4, HERC5, NUSAP1, SECISBP2) were totally missing, and 6 (DNAJC7, PITPNB, and RNF40, MTERF1, RBBP4, USP48) had only one quantitation, at 10 min; these were not analyzed further since their prediction were not robust. Significant proteins after this imputation were largely similar to those with no imputation, with some extra proteins increased in RNA association such as translation factor EIF3B; DExH-box helicase DHX15; and pre-mRNA maturation factor NUDT21, and some decreased in RNA association such as histone deacetylase HDAC1.

We conclude that imputation by Random Forest slightly increased numbers of proteins identified as significantly changed, while MINx0.5 made missing proteins appear significant.

### Comparing *in silico* and project-specific spectral libraries

All DIA analyses above were based on peptide search with spectral library constructed *in silico* in DIA-NN using light/heavy peptide fragments predicted from human proteome sequences obtained from Uniprot. We also tested the effects on protein quantitation of using a spectral library built on project-specific MS data. The total light proteome from HEK293T cells was purified, trypsin/Lys-C digested, C18 cleaned and acquired with four MS injections using DIA with different MS2 scan ranges between 400-1400 m/z. Total heavy spectra were calculated from the light data in DIA-NN and combined with the light spectra to form the library (detailed in Material and Methods). Comparisons between the DIA dataset analyzed with *in silico* and project-specific library are summarized in Fig. 5. Relative to the *in silico* dataset, the project-specific library detected 51 more proteins. However, 268 fewer proteins passed the threshold for analyses (detected in 2 out of 3 replicates at both time points in heavy control) and 224 fewer proteins were quantified at least twice at both time points. Of these, 34 more proteins passed the significance threshold. The project specific dataset had a lower maximum inter-replicate standard deviation than the *in silico* dataset for proteins with at least 2 quantitations at both time points (median=0.145 project-specific vs 0.164 *in silico*, compared to 0.197 for DDA dataset), indicating greater reproducibility.

**Fig. 5.**
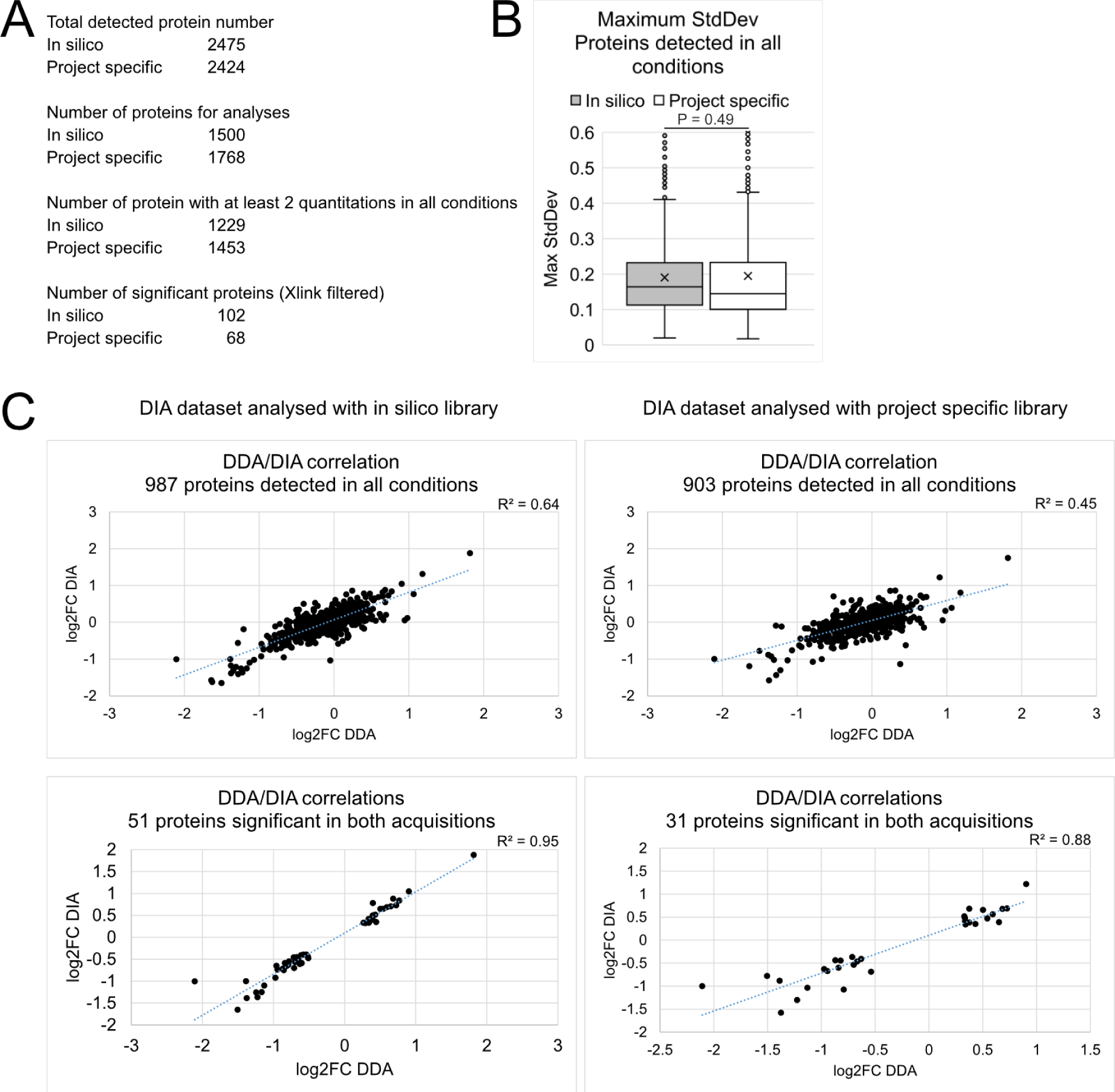
Effects of *in silico* or project-specific spectral libraries on DIA datasets. (A) Numbers of proteins: Totally detected; Selected for analysis (detected in at least 2 replicates at both time points in heavy controls); With at least 2 quantitations at each time points; Significantly altered plus enriched in UV-crosslinked samples when the peptides were searched using *in silico*-constructed or mass spec data-constructed (project-specific) spectral libraries. (B) Distribution of maximum inter-replicate standard deviations for proteins detected in all conditions in datasets generated with *in silico* or project-specific libraries. (C) DDA-DIA correlations for proteins detected in all conditions and significant in both acquisitions in datasets using *in silico* (left) or project-specific (right) libraries.

In terms of DDA-DIA correlation, the *in silico* reference dataset gave higher R^2^ values than the project-specific counterpart (0.64 vs 0.45 for proteins detected in all conditions). For the most similar proteins between DDA and DIA datasets, those classed as significantly altered in both, the R^2^ value was also higher in the *in silico* dataset (0.95 vs 0.88). It is possible that although the project-specific library had spectra that better matched the experimental samples, reflected by the lower inter-replicate variations indicating more consistent detections, the *in silico* library was constructed from the same proteomic sequences used in the DDA analysis and therefore had a more similar quantitation to DDA compared to the project-specific dataset. Proteins identified as significant in the *in silico* and project-specific datasets are listed in Table S2. It is noticeable that BH multiple test correction yielded only 3 significant proteins in the project-specific dataset, compared to 9 in the *in silico* dataset.

We conclude that peptide search using in silico spectral library produced better DDA-DIA correlation compared to a project-specific library. The *in silico* dataset was therefore chosen as the main dataset to compare with the DDA dataset in this study.

## Discussion

Quantitation of dynamic RNA-protein associations requires consistent relative measures of ratios between recovery of experimental samples and controls, which can be achieved using the SILAC labelling. However, conventional DDA-SILAC tandem-MS involves selective, and usually not customizable, peptide identification and quantitation, without MS2-level quality control. In contrast, DIA-SILAC does not apply selection between MS1 peptides, potentially quantifying a greater fraction of the total population, with precise precursor quality control at the MS2 level. This is reflected in the consistently lower inter-replicate variations in DIA datasets.

In the DDA datasets reported here, 1,069 proteins were reliably quantified between replicates, while the DIA datasets had 1,229 proteins - an increase of 15% over DDA. The two datasets identified overlapping genesets showing significant changes, enriched for translation regulators responding to cellular stress. However, the improved sensitivity in the DIA datasets allowed detection of potentially important proteins involved in arsenite stress response, including the transcription factor JUN. Together with the increased RNA association seen for its upstream signaling inhibitor PEBP1, this suggests a stress-responsive transcription regulation pathway, which is only visible in the DIA dataset.

Both datasets included many proteins that failed to be detected at one time point. These proteins would be classed as low abundance at these time points, and called as significantly increased/decreased. However, quantitation in other acquisitions frequently revealed quite different dynamics. Reliable quantitation therefore requires proteins to be detected at both time points, and the DIA dataset was superior in detecting such proteins. This was the case even without peptide fractionation following the pre-MS reverse phase column, and the consequential multiple MS injections per biological sample. DIA produced datasets that appeared more biologically informative, while reducing labor and machine time.

All methods of dataset normalization tested gave similar DDA/DIA correlations and inter-replicate variation. However, normalization by MaxQuant for DDA, and by Cyclic Loess for DIA gave the best balance between correlation and variation. The MINx0.5 imputation strategy called a unique set of significant proteins, mostly from those missing at 10 min.

While the Random Forrest approach called similar sets of significant proteins compared to no imputation, with 10 extra proteins. The two imputation strategies were based on different assumptions, with MINx0.5 assuming all missing proteins were due to low abundance and Random Forrest assuming they were missing at random. Therefore, the unique significant proteins identified by the two approaches may be useful in answering different questions.

The DIA raw files analyzed with a project-specific spectral library gave more consistent quantitation than using an *in silico* derive library for analysis, probably due to a better match with the experimental samples. In contrast, the *in silico* spectral library gave a better correlation between DIA and DDA datasets compared to the project-specific library. The DDA dataset was analyzed with the same proteomic sequence file used to generate the *in silico* library, probably allowing more similar quantitation between DDA and DIA.

We conclude that analyses of protein-RNA interactome using DIA can give superior results to the more conventional DDA approach, with reduced costs in labor and machine time.

## Supporting information

Non-crosslinked vs crosslinked cells

## Data availability

Original files for the datasets are available at EBI PRIDE database, https://www.ebi.ac.uk/pride/ (accession no. PXD045318).

## Acknowledgements

The authors thank Dr. Georg Kustatscher for valuable discussions in the concept and scientific content of this manuscript; Dr. Van Kelly for mass spec condition optimizations and help in confirming the DDA-DIA correlations at both peptide and protein levels; Dr. Vadim Demichev for development of DIA-NN and advice on DIA-SILAC analysis on GitHub; and Dr. Nic Robertson and Dr. Aleksandra Helwak for assistance with mammalian cell cultures.

## Supplementary data

**Fig.S1.**
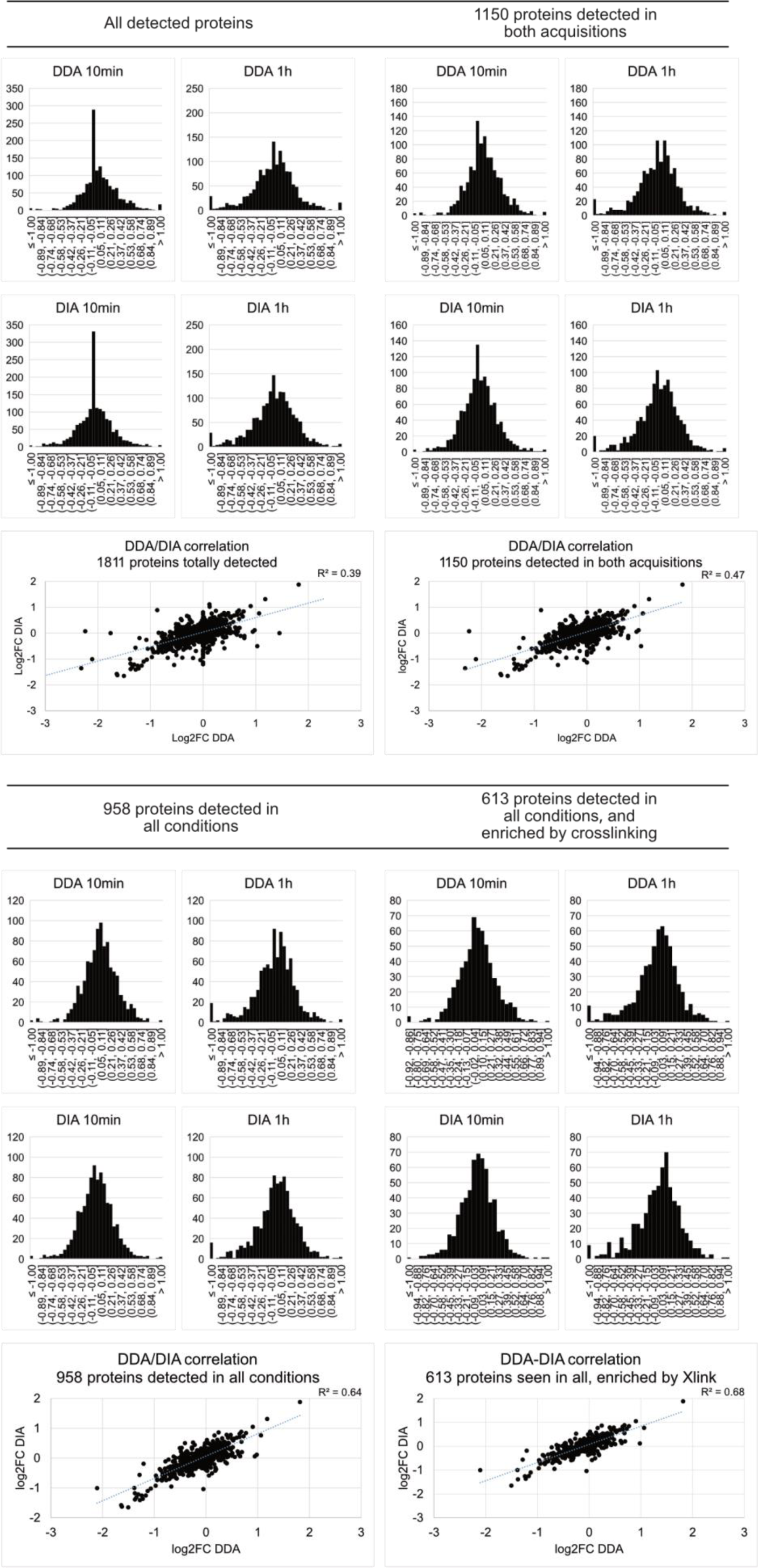
Distribution of Light:Heavy ratios and correlations of log2FC values between DDA and DIA datasets for 10 min and 1 h samples. Values were calculated from normalized, processed datasets. The DDA dataset was normalized by MaxQuant, DIA by Cyclic Loess. Separate plots are shown for; All detected proteins; Proteins detected in both acquisitions; Proteins detected in all conditions; Proteins detected in all conditions and enriched by UV-crosslinking.

**Fig.S2.**
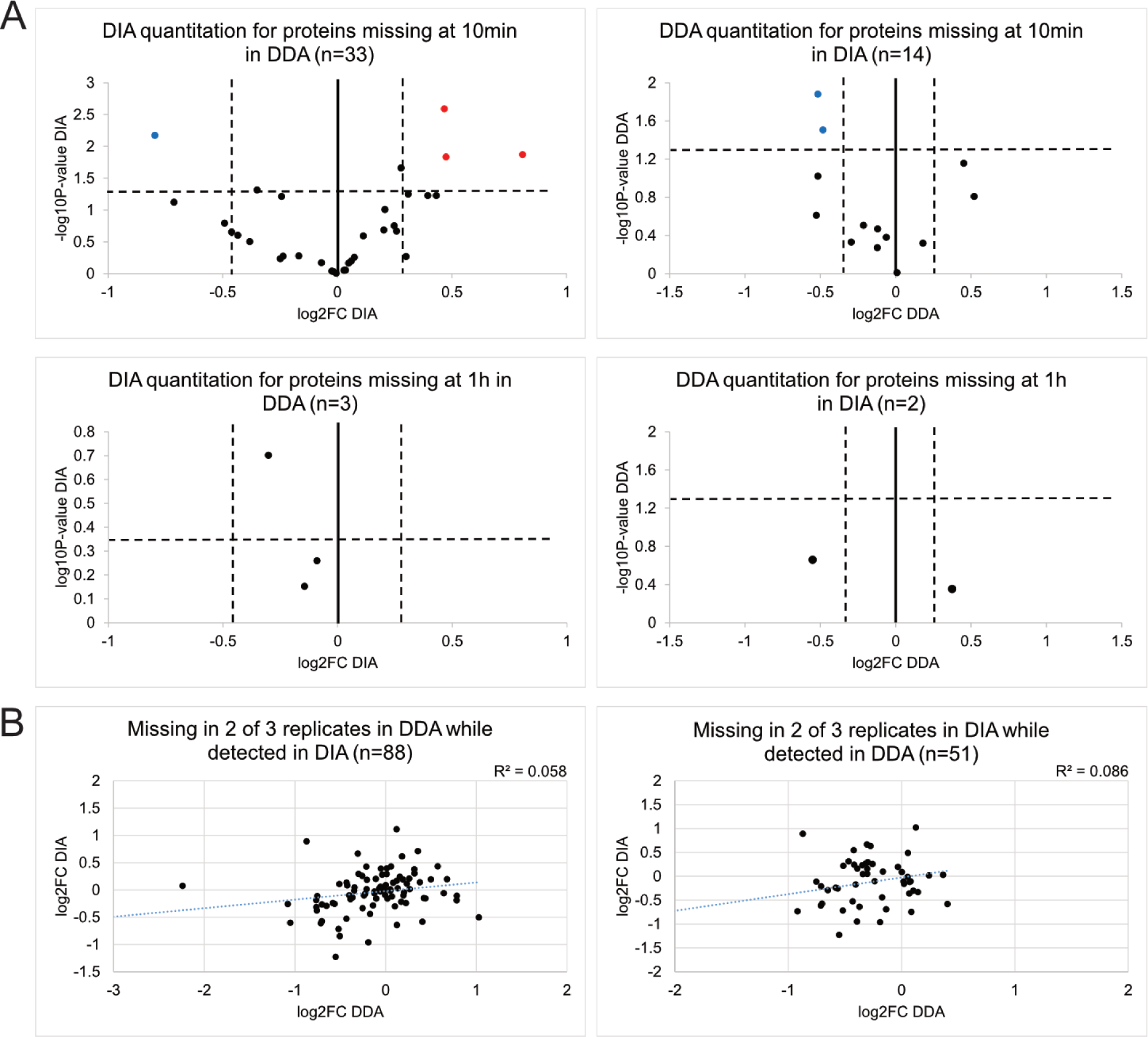
Proteins totally absent or with 2 missing values at any time point could not be consistently quantified between DDA and DIA datasets. (A) Volcano plot showing DDA or DIA quantitation for proteins totally missing at either 10 min or 1 h in the other acquisition. (B) DDA-DIA correlation for proteins with 2 missing values at any time point while also detected in the other acquisition.

**Fig.S3.**
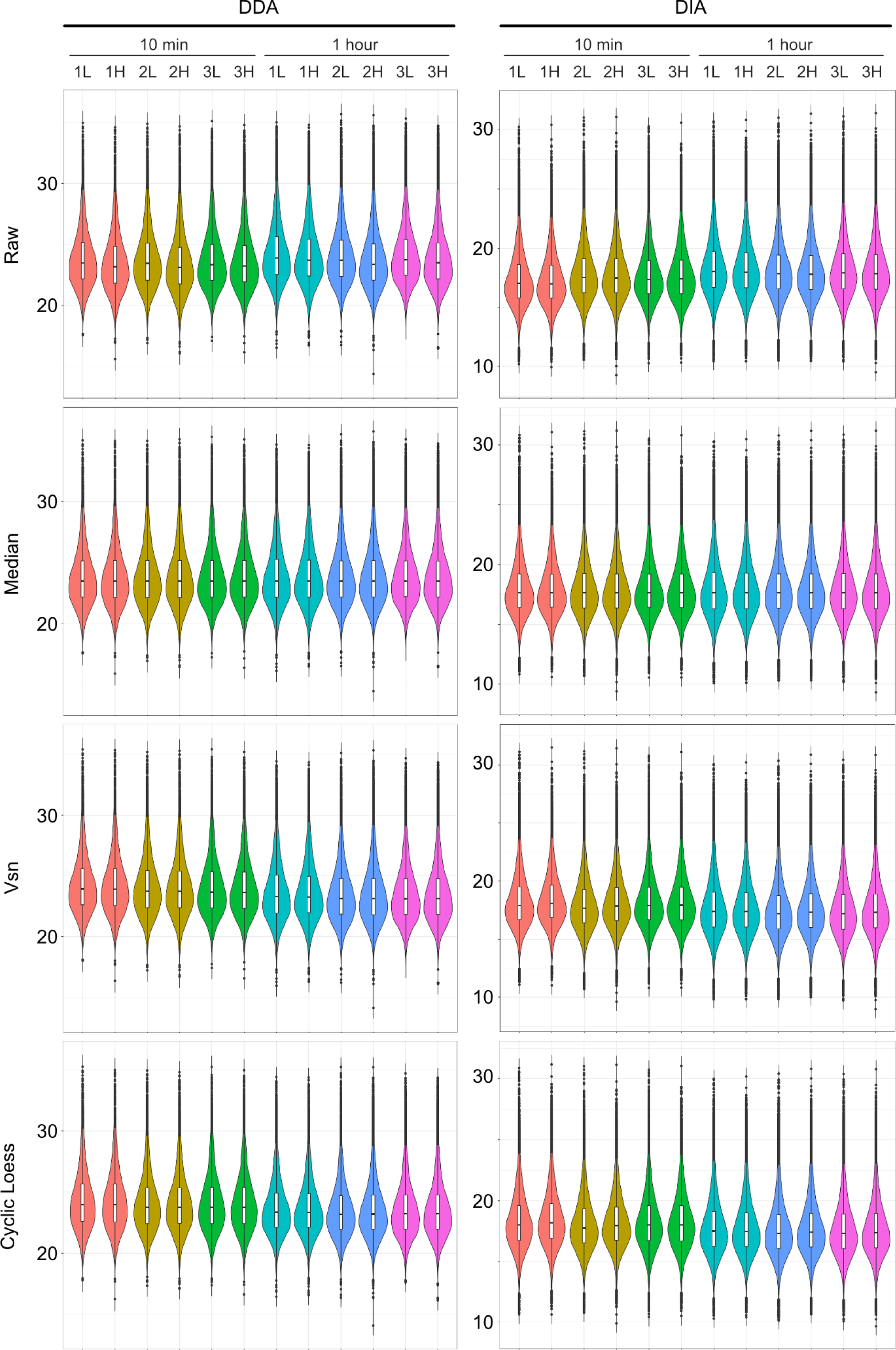
Effects of different normalization algorithms on distributions of peptide intensities. Raw intensities of individual precursors were normalized using median, Vsn, and Cyclic Loess in NormalyzerDE and plotted as violin and box plots. NA values were excluded. Center line, median; box limits, upper and lower quartiles; whiskers, 1.5x interquartile range; points, outliers.

**Fig.S4.**
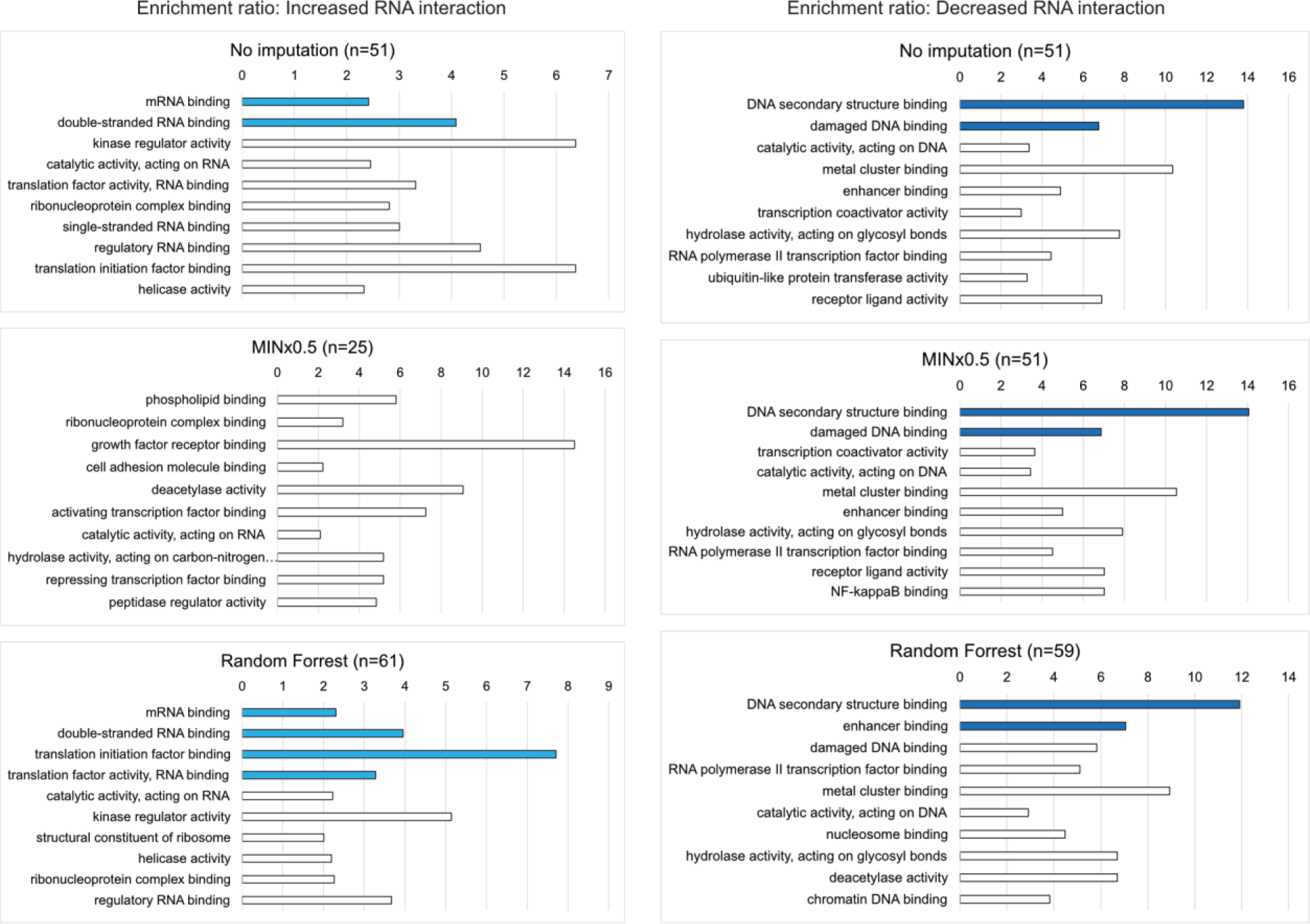
Top ten enriched GO terms for molecular function in datasets with: No imputation; Imputation by MINx0.5; Imputation by Random Forrest. GO term enrichment was analyzed for proteins with significantly increased (left) or decreased (right) RNA interactions from 10 min and1 h samples in the DIA datasets. These were selected using 2-sided T Test: P value <0.05 and fold-change >90^th^ and <10^th^ percentile. Dark blue bar, FDR<0.05; pale blue bar, FDR between 0.05-0.06; white bar, FDR>0.05. GO terms were sorted by FDR in ascending order, or by enrichment ratio if all FDRs were 1.

**Table S1.** List of proteins showing significant changes in the three datasets. Dark grey shaded, consistent between all three; Pale grey shaded, consistent between no imputation and Random Forrest; Bold black font, proteins significant after Benjamini-Hochberg multiple test correction, FDR<0.05.

| Increase RNA Interaction |  |  | Decrease RNA Interaction |  |  |
| --- | --- | --- | --- | --- | --- |
| No imputation | MINx0.5 | Random Forrest | No imputation | MINx0.5 | Random Forrest |
| ATXN2 | **DBNL** | ATXN2 | ATAD5 | **APRT** | ATAD5 |
| BMS1 | DNAJC7 | BMS1 | BAZ1A |  |  |
| CCDC9 | **EIF2A** | CCDC9 | BAZ1B | BAZ1B | BAZ1B |
| CPEB4 | **MBNL3** | CPEB4 | BAZ2A | BAZ2A | BAZ2A |
| CSTF1 | **MRPL22** | CSTF1 | BCOR | BCOR | BCOR |
| DCAF13 | **MRPS25** | **DCAF13** | CCDC124 | CCDC124 | CCDC124 |
| DDX3X | **MRPS35** | DDX3X | CCDC86 | CCDC86 | CCDC86 |
| DDX54 | **NMD3** | DDX54 |  | **CDKN2A** | **CDCA2** |
| DDX6 |  | DDX6 | CHAF1A | CHAF1A | CHAF1A |
|  |  | DHX15 | CHAMP1 | CHAMP1 | CHAMP1 |
| **DHX9** |  | **DHX9** |  | **COPZ1** | **CNOT4** |
| DKC1 |  | DKC1 | DRG1 | DRG1 | DRG1 |
|  |  | DNAJC7 | EDF1 | **EDF1** | **EDF1** |
| EIF3A |  | **EIF3A** | FCF1 | FCF1 | FCF1 |
|  |  | EIF3B | G3BP1 | G3BP1 | G3BP1 |
| EIF4A1 |  | EIF4A1 | GNL3L | GNL3L | GNL3L |
| EIF4B |  | EIF4B | GTF3C1 | GTF3C1 | GTF3C1 |
| EIF4H |  | EIF4H |  |  | HDAC1 |
| FXR2 |  | FXR2 | HDLBP | HDLBP | HDLBP |
|  |  | IMPDH2 |  |  | HERC5 |
| IQGAP1 |  | IQGAP1 | HMGB1 | HMGB1 | HMGB1 |
| KRR1 |  | KRR1 | HMGB2 | HMGB2 | HMGB2 |
| MFAP1 |  | MFAP1 | HMGB3 | HMGB3 | HMGB3 |
| MRPL10 |  | MRPL10 | **HNRNPA0** | **HNRNPA0** | **HNRNPA0** |
| MRPL2 |  | MRPL2 | INCENP | INCENP | INCENP |
| MRPL34 |  | MRPL34 | JUN | JUN | JUN |
| NOB1 |  | NOB1 | KDM3B | **KDM3B** | KDM3B |
|  |  | NUDT21 |  |  | KIF22 |
| PARP12 |  | PARP12 | LIG3 | LIG3 | LIG3 |
| PATL1 |  | PATL1 | LTV1 | LTV1 | LTV1 |
| PEBP1 | PEBP1 | PEBP1 | MGA | **MGA** | MGA |
| PES1 | **PHAX** | PES1 | MKI67 | MKI67 | MKI67 |
|  | **PRCC** | PITPNB | MPG | MPG | MPG |
| PRPF8 | RNF214 | **PRPF8** | MTDH | MTDH | MTDH |
| PRR3 | **RPP25L** | PRR3 |  | NEDD8 | **MTERF1** |
| PRRC2A | **RPP30** | PRRC2A |  |  | **NUSAP1** |
| PRRC2C | SAMD4B | PRRC2C | PDCD4 | PDCD4 | PDCD4 |
| PUM1 | **SIN3A** | PUM1 | POLE | POLE | POLE |
| PUM2 | **SMNDC1** | PUM2 | PSMC3 | PQBP1 | PSMC3 |
| R3HDM1 | **SNX2** | R3HDM1 | RAI1 | RAI1 | RAI1 |
| R3HDM2 | **SUPV3L1** | R3HDM2 |  |  | **RBBP4** |
| RBFOX2 | **THUMPD1** | RBFOX2 | RBM42 | RBM42 | RBM42 |
| RBM19 | **TRNAU1AP** | RBM19 | RIF1 | RIF1 | RIF1 |
| RBM8A | TUT1 | RBM8A | RPL11 |  |  |
|  | **YTHDF3** | RNF40 | RPS10 | RPS10 | RPS10 |
| RPA1 | ZC3H15 | RPA1 | **RPS14** | **RPS14** | **RPS14** |
| RPL17 | **ZC3H6** | RPL17 | **RPS28** | **RPS28** | **RPS28** |
|  |  | RPL18A | RPS3 | **RPS3** | RPS3 |
| RPL29 |  | RPL29 | RPS5 | **RPS5** | RPS5 |
| RPL35 |  | RPL35 |  |  | **SECISBP2** |
|  |  | RPS16 |  |  | **SLC4A1AP** |
| RPS24 |  | RPS24 | **SND1** | **SND1** | **SND1** |
| **SART3** |  | **SART3** | SRP68 | SRP68 | SRP68 |
| TRIM25 |  | TRIM25 | TARDBP | TARDBP | TARDBP |
| UTP15 |  | UTP15 | TMPO | **TMPO** | **TMPO** |
| WDR75 |  | WDR75 | TRIP12 | **UBE2N** | TTF2 |
| WNK1 |  | WNK1 | UBE2D2 |  |  |
| YTHDC2 |  | YTHDC2 | UHRF1 | UHRF1 | UHRF1 |
| ZC3HAV1 |  | ZC3HAV1 | **UPF1** | **UPF1** | **UPF1** |
|  |  | ZCCHC3 |  |  | **USP48** |
| ZNF622 |  | ZNF622 | **XPC** | **XPC** | **XPC** |

**Table S2.** List of proteins showing significant changes in DIA datasets analyzed with *in silico* or project-specific spectral libraries. Level of similarity between the two analyses are indicated above each list. Significance threshold: 2-sided T Test P-value <0.05, FC >90^th^ and <10^th^ percentile. Proteins shown in bold font indicate those deemed significant after Benjamini Hochberg multiple test correction (FDR<0.05).

|  |  |
| --- | --- |
| ATXN2 | ATXN2 |
| BMS1 |  |
| CCDC9 | CCDC9 |
| CPEB4 |  |
| CSTF1 |  |
| DCAF13 |  |
| DDX3X | DDX3X |
| DDX54 | DDX54 |
| DDX6 |  |
| **DHX9** | DHX9 |
| DKC1 |  |
| EIF3A | EIF3A |
| EIF4A1 | EIF4A1 |
| EIF4B | EIF4B |
|  | EIF4G2 |
| EIF4H | **EIF4H** |
|  | FMR1 |
| FXR2 | FXR2 |
| IQGAP1 |  |
| KRR1 | KRR1 |
| MFAP1 | LSM6 |
| MRPL10 | LSM8 |
| MRPL2 | METTL16 |
| MRPL34 | MRPS2 |
| NOB1 | NOC3L |
| PARP12 | NUDT21 |
| PATL1 | PPAN |
| PEBP1 |  |
| PES1 |  |
| PRPF8 |  |
| PRR3 | PRR3 |
| PRRC2A |  |
| PRRC2C | PRRC2C |
| PUM1 | PUM1 |
| PUM2 | PUM2 |
| R3HDM1 | **R3HDM1** |
| R3HDM2 | R3HDM2 |
| RBFOX2 | RBM34 |
| RBM19 | RBMXL1 |
| RBM8A |  |
| RPA1 | RPA1 |
| RPL17 | RPL17 |
| RPL29 | RPL3 |
| RPL35 | RPS2 |
| RPS24 | SAFB |
| **SART3** | TRUB1 |
| TRIM25 | UPF3B |
| UTP15 |  |
| WDR75 |  |
| WNK1 |  |
| YTHDC2 |  |
| ZC3HAV1 |  |
| ZNF622 |  |

**Table S2. List of proteins showing significant changes in DIA datasets analyzed with *in silico* or project-specific spectral libraries.**
|  |  |
| --- | --- |
| ATAD5 |  |
| BAZ1A |  |
| BAZ1B | BAZ1B |
| BAZ2A |  |
| BCOR | BCOR |
| CCDC124 |  |
| CCDC86 |  |
| CHAF1A | CHAF1A |
| CHAMP1 | CHAMP1 |
| DRG1 | CHTF18 |
|  | CKAP4 |
|  | CNOT4 |
| EDF1 | EDF1 |
| FCF1 | EMD |
| G3BP1 | FNDC3A |
| GNL3L |  |
| GTF3C1 | GTF3C1 |
| HDLBP | HP1BP3 |
| HMGB1 |  |
| HMGB2 |  |
| HMGB3 |  |
| **HNRNPA0** |  |
| INCENP |  |
| JUN |  |
| KDM3B |  |
| LIG3 | LIG3 |
| LTV1 | LTV1 |
|  | MECP2 |
| MGA | MGA |
| MKI67 | MKI67 |
| MPG | MPG |
| MTDH | MTDH |
| PDCD4 |  |
| POLE | POLE |
| PSMC3 | RBBP4 |
| RAI1 | RBM5 |
| RBM42 | RBM7 |
| RIF1 |  |
| RPL11 | RPL11 |
| RPS10 |  |
| **RPS14** | RPS14 |
| **RPS28** | RPS28 |
| RPS3 | RPS3 |
| RPS5 | RPS5 |
|  | SBDS |
|  | SLC4A1AP |
| **SND1** | **SND1** |
| SRP68 |  |
| TARDBP |  |
| TMPO | TMPO |
| TRIP12 |  |
| UBE2D2 |  |
| UHRF1 | UHRF1 |
| **UPF1** |  |
| **XPC** |  |
| **ZC3H11A** |  |

